# A novel gammaproteobacterial methanotroph from *Methylococccaeae*; strain FWC3, isolated from canal sediment from Western India

**DOI:** 10.1101/696807

**Authors:** Kumal Khatri, Pranitha S. Pandit, Jyoti Mohite, Rahul A. Bahulikar, Monali C. Rahalkar

## Abstract

We isolated a gammaproteobacterial methanotrophic strain FWC3, from canal sediment from Western India. The strain oxidizes methane and can also grow on methanol. The draft genome of the same was sequenced which showed a size of ∼3.4 Mbp and 63% GC content. FWC3 is a coccoid, pale pink pigmented methanotroph and is seen in the form of diplococci, triplets, tetrads or small aggregates. After comparison of the complete 16S rRNA gene sequence, average amino-acid similarities and digital DNA-DNA hybridization values with that of the neighboring type species, we propose that the strain belongs to a novel genus and species, *Ca*. Methylolobus aquaticus FWC3^Ts^.

---

Aerobic methanotrophs play a very important role in mitigation of methane, the second most important greenhouse gas (1). Strain FWC3, a *Methylococcaceae* member, was isolated (in 2018) from a fresh-brackish water canal sediment sample collected from Nagaon, Alibag, India (18.65° N; 72.86° E) in 2017, using enrichment and isolation method as described (2). Recently, we described isolation, cultivation of several novel members of *Methylococcaceae* from Indian wetland habitats and reported their draft genomes (3-7) and FWC3 is a new addition. The strain forms pink colonies on agarose medium incubated with methane in the atmosphere (Fig.1 a) and oxidizes methane when grown in liquid medium with methane and air in gas phase. FWC3 could also tolerate and grow in methanol containing medium up to 4% concentrations. Strain FWC3 shows coccoid to slightly oval-shaped cells with a diameter of ∼1-1.5 µm and the cells are present as diplococci, triplets, tetrad or small groups (Fig. 1b).

**Fig. 1.**
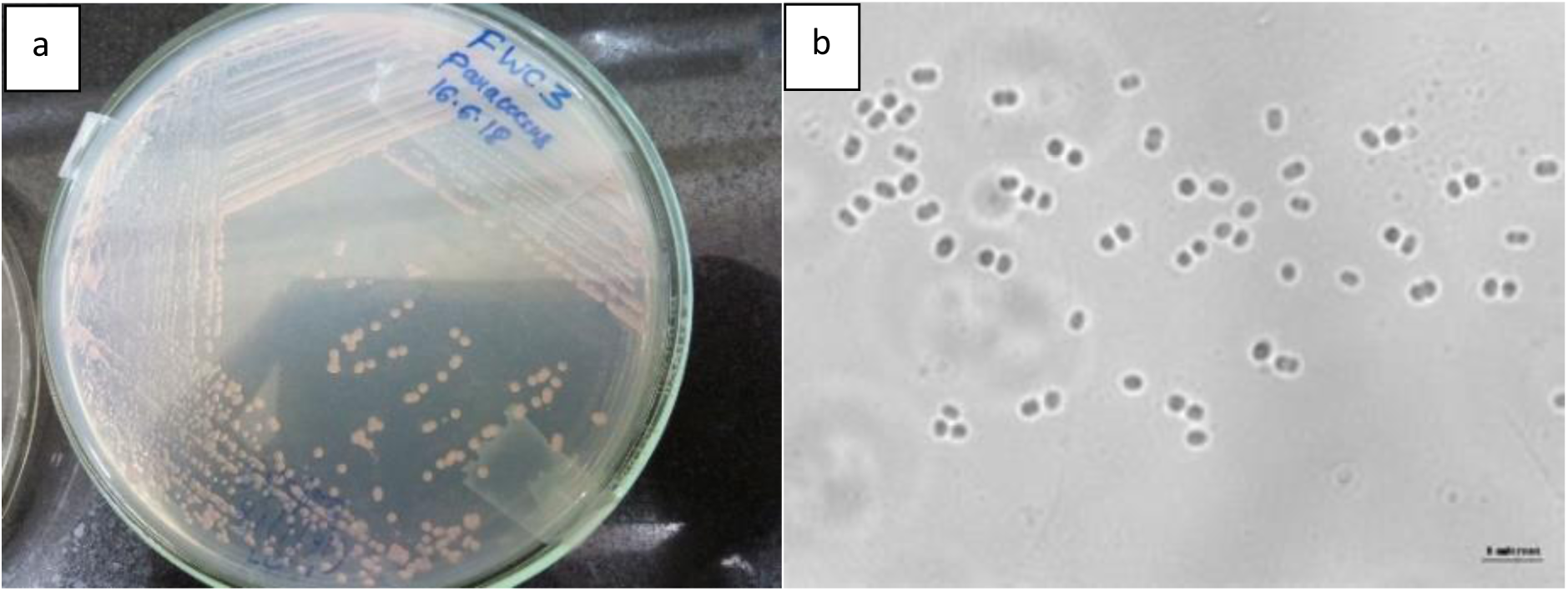
a. FWC3 growing on dilute NMS agarose plates in 20% methane environment. 1b. Phase contrast image of FWC3 growing on methane, the bar represents 5 µm.

Partial 16S rRNA gene sequence (∼1490 bp) of FWC3 showed only 94.3% to *Methylocaldum marinum* S8^T^ and lesser similarity with other *Methylococcaceae* members. The genome of FWC3 was sequenced for understanding the taxonomic novelty and metabolic properties.

Genomic DNA of FWC3 was extracted from an axenic culture, grown on solid medium, using Gen Elute™ Bacterial Genomic DNA kit, Sigma-Aldrich. The genome was sequenced using Illumina HiSeq platform (2 × 150 bp) in Medgenome laboratories, Bangalore. A minimum of 75% of the sequenced bases were of Q30 value. The sequenced data was processed to generate FASTQ files and uploaded on the FTP server for download and secondary analysis. A total of 16 million paired end reads which were filtered for low quality bases and adapter sequences using Trimmomatic (*ν 0*.*35*) and cutadapt *(ν 1*.*18*) to retain high quality reads. Approximately 8 million high quality paired end reads were used for assembly with SPADES (*ν 3*.*13*.*0*) assembler. The size of FWC3 draft genome is 3.4 Mbp with a GC content of 63%. A total of 42 scaffolds of >500 bp were constructed, with an N50 of 184 kb, the largest scaffold assembled measured 498.3 kb and the total coverage was 700X. Annotation was performed in Rapid Annotation using Subsystem Technology (8) (RAST), and annotation from the NCBI prokaryotic genome annotation pipeline. In total, 2831 proteins were annotated as per NCBI annotation and similar values were obtained using RAST. The whole genome shotgun project is open to public access since February 2019. The whole genome is in accordance with the minimal standards specified for the use of taxonomic data for prokaryotes with 600-700X coverage (9). When the genome of FWC3 was compared with the nearest neighboring methanotrophs *Methylocaldum marinum* S8 and *Methylococcus capsulatus* Bath using the average amino-acid identities (AAI). The AAI values were 59.2% and 60.7%, respectively and the average nucleotide indices were ∼70% well below the genome cut-off. The digital DNA-DNA hybridization of FWC3 strain with *Methylocalcum marinum* S8 and *Methylococcus capsulatus* Bath are 19% +/-2 respectively. Thus, FWC3 could be proposed to be a member of a novel genus based on the whole genome data.

The cell wall fatty acids of strain FWC3 showed maximum amounts of 16:1 ω6c/ 16:1 ω7c (41%) followed by 16:0 (26%) and 16:1 ω9c (16%) and hence a unique pattern was observed, compared to other Type Ib methanotrophs.

Additionally, the complete 16S rRNA sequence derived from the genome assembly MN080433.1 (1527 bp), shows 94.39% similarity to *Methylocaldum marinum* S8^T^ based on the blast analysis done on 09.07.2019. A 1000 maximum likelihood bootstrap phylogenetic tree constructed also shows the prominent position of strain FWC3 branching with other Type Ib methanotrophs but forming a distinct branch (Fig. 2).

**Fig. 2.**
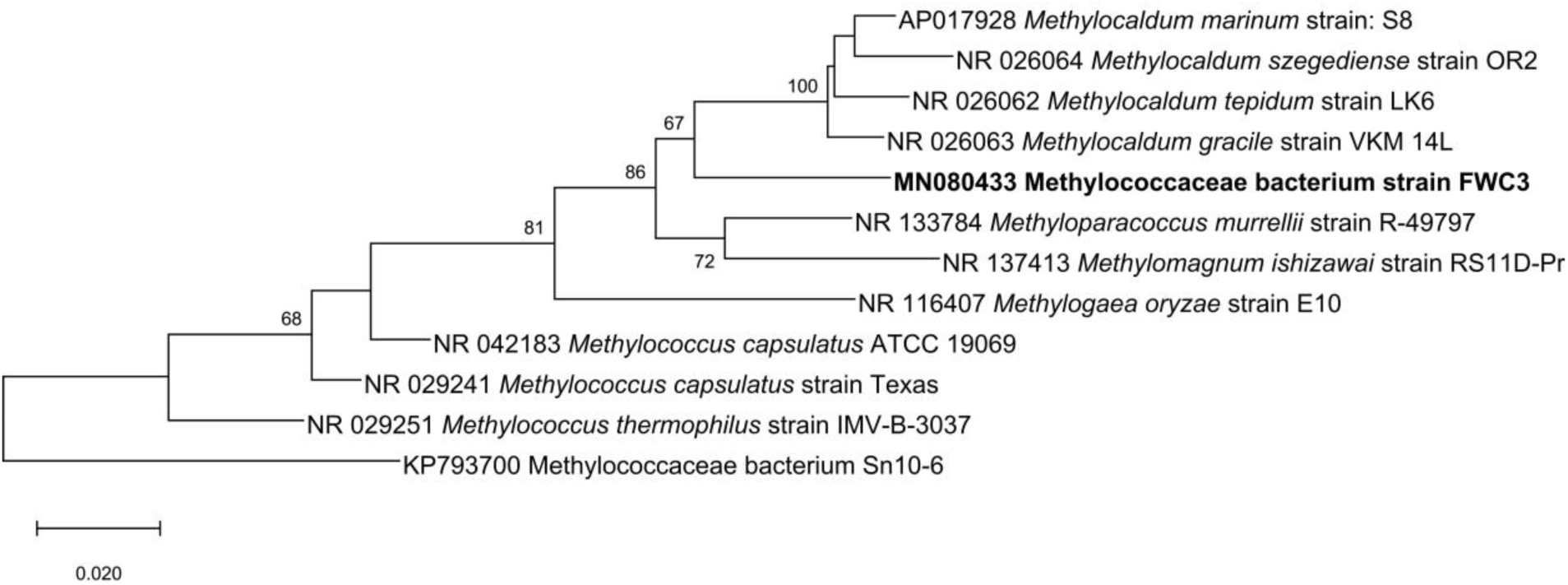
Maximum likelihood tree using 1000 bootstraps showing the phylogenetic position of the 16S rRNA gene of FWC3 (MN080433) with respect to other TypeIb methanotroph type species. The bar represents 2% sequence distance. The tree is based on the sequences appearing in the NCBI nucleotide blast as on 9^th^ July 2019. The tree was constructed using MEGA 6.0 (10).

At present, FWC3 cannot be grown to a sufficient extent and preserved for long term storage, which are requirements for the preservation in two international foreign culture collections located in two different countries. Hence, we propose a Candidatus name, ‘*Candidatus* Methylolobus aquaticus’ FWC3 ^Ts^ (taxonomic novelty based on the whole genome sequence). This proposal is as per the current proposal for describing novel species based on the genome sequence and due to the fact that our genome fulfills the minimal standards for the use of genome data for taxonomic purposes (9, 11).

## Database availability

The whole genome shotgun project has been deposited in GenBank database under the accession number <u>NZ_SEYW00000000.1</u> (released in Feb 2019) under the name, *Methylococcaceae* bacterium FWC3. The bioproject accession number is PRJNA52097. The link for the genome is https://www.ncbi.nlm.nih.gov/genome/?term=SEYW01.

## Acknowledgments

We acknowledge MACS Agharkar Research institute, MCR acknowledges Department of Science and Technology, DST, SERB (EMR/2017/002817) and PSP acknowledges DST (SR/WOS-A/LS-410/2017) for providing the funds. Kumal Khatri acknowledges CSIR for providing her fellowship.

